# Accessible, Reproducible, and Scalable Machine Learning for Biomedicine

**DOI:** 10.1101/2020.06.25.172445

**Authors:** Qiang Gu, Anup Kumar, Simon Bray, Allison Creason, Alireza Khanteymoori, Vahid Jalili, Björn Grüning, Jeremy Goecks

## Abstract

Supervised machine learning, where the goal is to predict labels of new instances by training on labeled data, has become an essential tool in biomedical data analysis. To make supervised machine learning more accessible to biomedical scientists, we have developed Galaxy-ML, a platform that enables scientists to perform end-to-end reproducible machine learning analyses at large scale using only a web browser. Galaxy-ML extends Galaxy, a biomedical computational workbench used by tens of thousands of scientists across the world, with a machine learning tool suite that supports end-to-end analysis.

---

Machine learning (ML) has become an essential tool in biomedicine to make sense of large, high-dimensional datasets such as those found in genomics, proteomics, and imaging^1–3^. In supervised machine learning, these datasets are used to build statistical models from high-dimensional feature sets that can predict continuous values (regression analysis) or discrete classes (classification). Example applications of ML to biomedicine include developing predictive models for drug metabolism rates using brain images^4,5^, genotype-phenotype associations^3^, and drug response in model systems^6,7^. Deep learning, which leverages multi-layer neural networks, has been used for prediction of splice sites^8^, protein structures^9^, and the presence of cancer from histopathology images^10^.

Many software libraries, tools, and platforms have been developed to make machine learning easier to use. Software libraries such as scikit-learn^11^ and Keras^27^ provide programmatic abstractions that make it simpler to build machine learning pipelines for bioinformaticians or computational scientists. Selene^12^ and Ludwig (https://github.com/uber/ludwig) use higher-level abstractions in the form of configuration files and predefined building blocks like encoders and neural network architectures to simplify deep learning. Neural network visualization and development toolkits such as Tensorboard (https://www.tensorflow.org/guide/summaries_and_tensorboard) and Azure ML Studio (https://studio.azureml.net/) provide interactive interfaces for understanding and improving neural networks.

Despite these successes, machine learning is often difficult to use in biomedicine. A successful application of machine learning to biomedical data must be able to integrate multiple analysis tools, be easily accessible, scale to large datasets, and reproduce results. Most ML tools lack a graphical user interface, reducing accessibility for scientists with limited informatics skills. As the size and number of biomedical datasets continues to grow, computational infrastructure such as workflow engines and job schedulers are needed for scaling and reproducing machine learning applications in biomedicine and for benchmarking multiple machine learning methods. Addressing these challenges requires an integrated software solution that (1) makes machine learning accessible to biomedical scientists who have limited programming and informatics knowledge and (2) connects machine learning with the broader ecosystems of biomedical analysis tools and a scalable computational workbench.

To meet this need, we have developed Galaxy-ML (Figure 1), an extension of the Galaxy platform (http://galaxyproject.org)^13^ that features a large and diverse suite of supervised machine learning tools. Galaxy is a user-friendly web-based computational workbench used by tens of thousands of scientists across the world for a wide variety of biomedical data analysis, including genomics, proteomics, metabolomics, cheminformatics, image processing, and flow cytometry. A key aspect of Galaxy is its web-based user interface, enabling anyone to use complex analysis tools and multi-tool workflows without any knowledge of programming. Galaxy-ML uses the Galaxy web interface to make machine learning tools and pipelines widely accessible.

**Figure 1.**
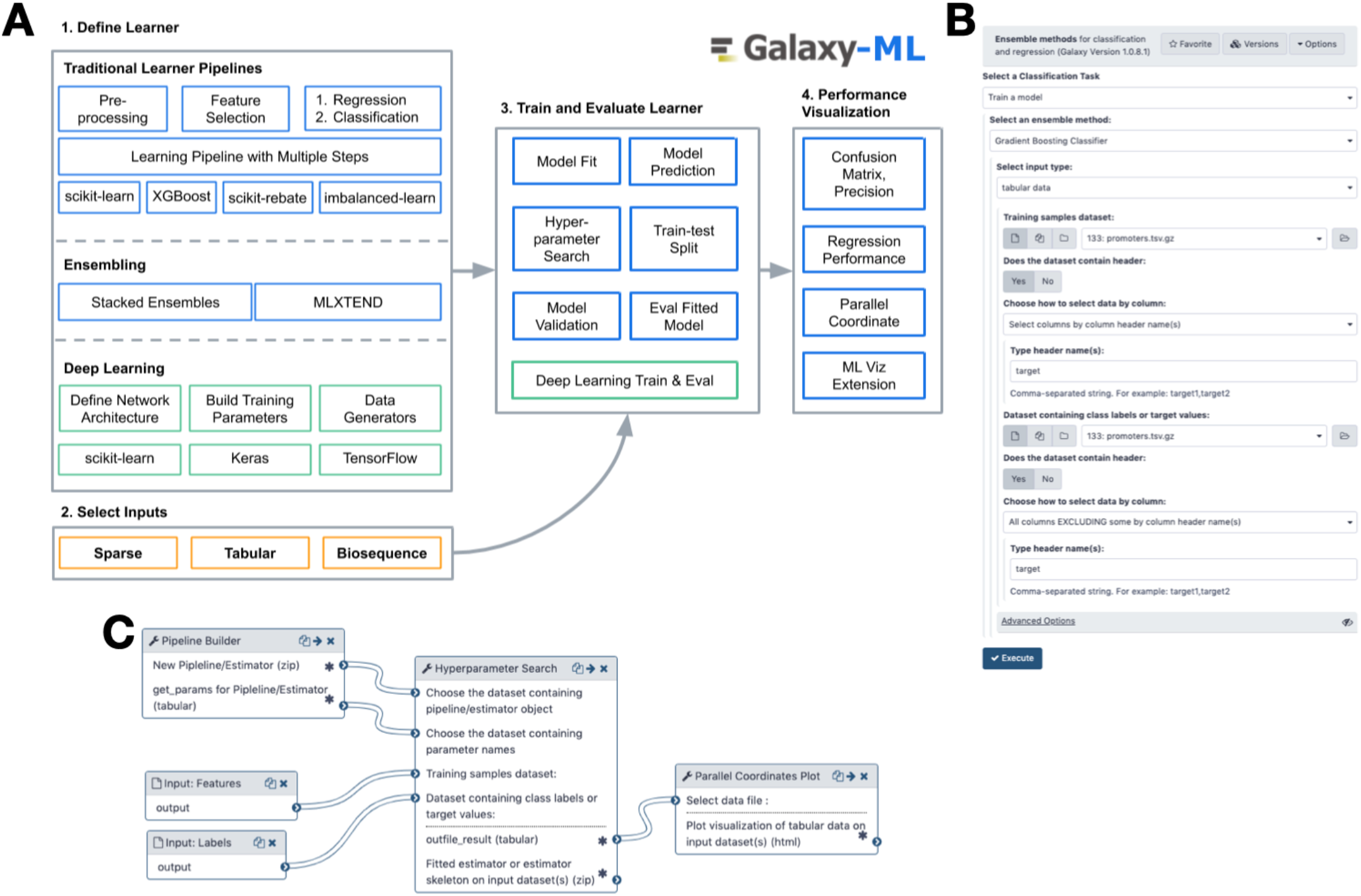
Panel A: The Galaxy-ML platform provides all the tools necessary to define a learner, train it, evaluate it, and visualize its performance. Panel B is a screenshot of the Galaxy tool to create a gradient boosted classifier. Panel C shows a Galaxy workflow to create a learner using a pipeline, perform hyperparameter search, and visualize the results.

Galaxy-ML also provides benefits in scalability, reproducibility, and workflow development. Large machine learning analyses, such as optimising hyperparameters and model evaluation across many different datasets, can require executing tens of thousands of analyses. Galaxy-ML uses Galaxy’s workflow system to execute large-scale analyses by distributing them across one or more computing clusters and running them in parallel. Galaxy ensures reproducibility by recording all parameters and tools used, so all analyses, including those for machine learning, are completely reproducible. This is critical, as reproducibility continues to gain importance in machine learning research^14^. Galaxy-ML enables end-to-end machine learning analyses that begin with processing primary biological data and end with trained machine learning models that can make predictions of phenotypic attributes like demographics or prognosis. For instance, a tutorial^15^ is provided in which Galaxy-ML is utilized to reproduce a study that uses RNA-seq data to predict an individual’s chronological age^16,17^. End-to-end workflows are possible because Galaxy-ML’s machine learning tools can be connected to the more than 7,800 tools available in the Galaxy ToolShed^18^ for analyzing genomics, proteomics, imaging, and other kinds of biomedical data.

Galaxy-ML supports four major steps in machine learning—preprocessing, modeling, ensembling, and evaluation—by integrating six machine learning libraries (Table 1) together with additional visualization and conversion tools. Scikit-learn^11^ provides the foundation for Galaxy-ML with approaches for all four major steps; most scikit-learn methods are available in Galaxy-ML. Additional libraries are included to meet key needs for machine learning in biomedicine, including feature selection, approaches for working with imbalanced datasets, and modeling approaches using gradient boosted decision trees, deep learning, and ensembling. Using Galaxy-ML, tools from all these libraries can be connected together into complete machine learning pipelines and can be stored and reused as an executable workflow. An example pipeline for using RNA-seq to predict drug response will (1) normalize gene expression values to bring them into the same scale; (2) select genes that show the highest variance; (3) use the selected genes to create a predictive model such as logistic regression or gradient-boosted decision trees; (4) use grid search with cross validation to optimize model hyperparameters; and (5) visualize model performance from the grid search using a heatmap or parallel coordinates plot. Galaxy-ML can be used to create thousands of different machine learning pipelines. Documentation, along with tutorials, is available at https://galaxyproject.org/community/machine-learning/, and links to the Galaxy-ML code and tool repositories are available in the Methods section.

**Table 1.**
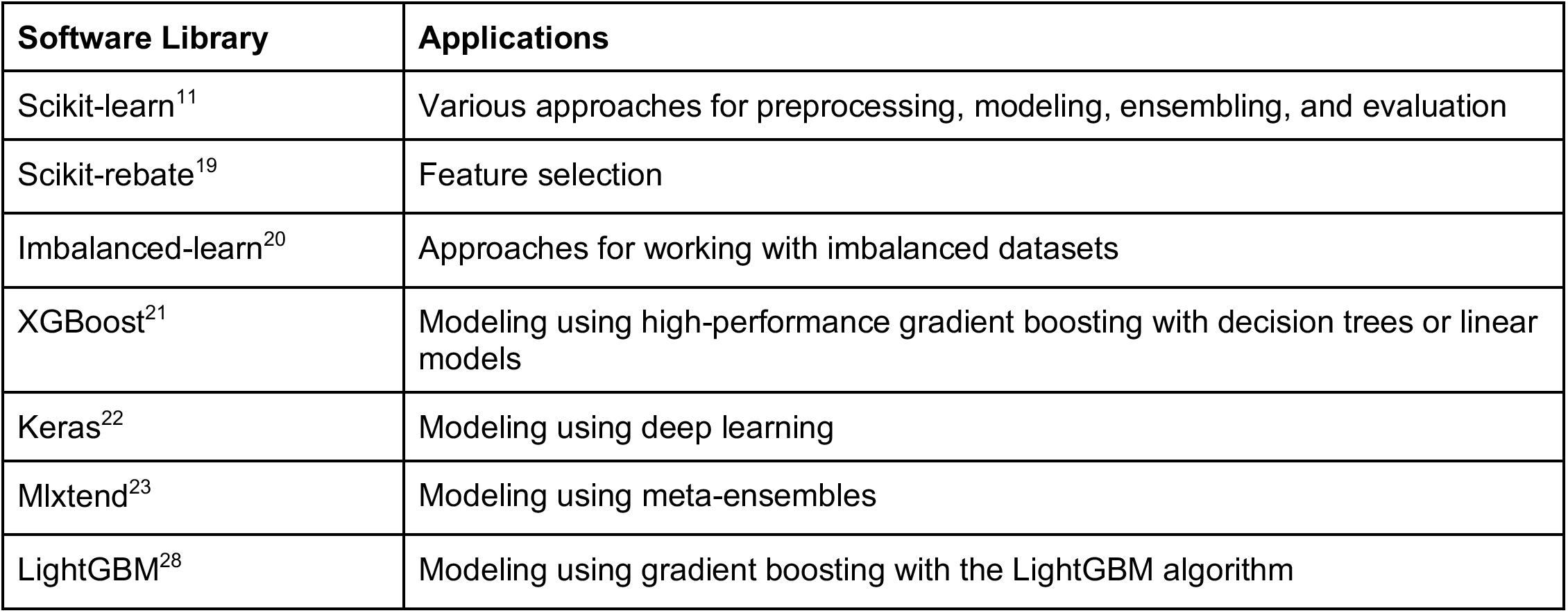
Software libraries integrated into Galaxy-ML and their applications.

We demonstrate the utility of Galaxy-ML in three use cases: (1) extending a machine learning benchmark experiment where 4,000 models were created and evaluated on 276 biomedical datasets^24^; (2) predicting drug response activity in cancer cell lines using gene expression datasets using stacked meta-ensembles; and (3) recreating deep learning models for genomics that predict, among other attributes, the functional impact of genetic variants. The Methods section provides links to complete analysis histories and results so that all analyses can be fully reproduced on any Galaxy server with the Galaxy-ML tool suite. All analyses were performed on a public Galaxy server at https://usegalaxy.eu and are listed at https://ml.usegalaxy.eu. All workflows, data and results can be accessed via a web browser and analyses can be reproduced directly.

In the first use case, we used Galaxy-ML to extend an analysis of machine learning models across 276 biomedical datasets^24^—164 classification datasets and 112 regression datasets^25^. The original analysis compared performance of 13 models on the 164 classification datasets. We applied 15 models to the classification datasets and 14 models to the regression datasets, creating a total of 4,028 trained models with hyperparameters optimized using grid search. We evaluated all models using 10-fold cross-validation (CV). Because many datasets were imbalanced, *F1* scoring rather than ROC AUC was used to evaluate performance of classification models, and Pearson’s R^2^ was used to evaluate performance of regression models. Performance of classification models are concordant with the initial publication: (a) boosted tree models perform best overall (Figure 2a) and (b) automated hyperparameter optimization improves performance for many models (Figure 2b). Performance of regression models are similar to those in classification, though boosted tree models only modestly outperform random tree models, and hyperparameter optimization often improves results most for models with low overall performance (see Methods section M.1.1 for more details).

**Figure 2.**
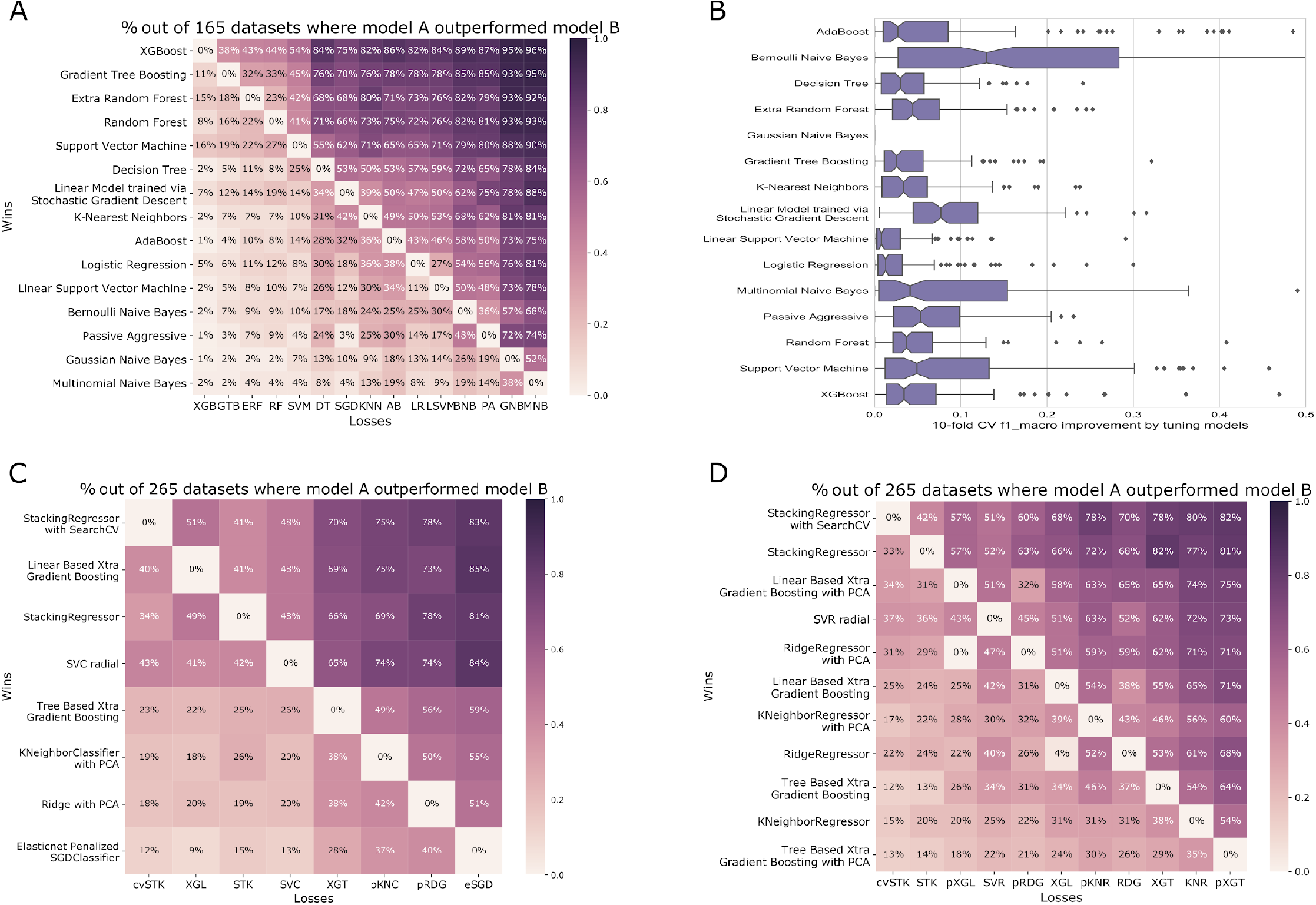
Pairwise performance comparisons for use cases 1 and 2. Use case 1 pairwise comparisons for classification tasks on 164 structured biomedical datasets^24^ show decision tree forests perform best (panel A) and hyperparameter optimization can improve the performance of most models (panel B). Use case 2 results for prediction using regression (panel C) and classification (panel D) show ensemble approaches that use stacking perform best, though linear-based gradient boosting also performs. In panels A, C, and D, heatmaps show the percentage of datasets for which the model listed along the row outperforms the model along the column. For instance, in panel A, XGBoost outperforms Gradient Tree Boosting (GTB) from scikit-learn on 38% of datasets, GTB outperforms XGBoost on 11% of datasets, and they perform equivalently on 51% of datasets.

For the second use case, we implemented stacked meta-ensemble predictors in Galaxy-ML for drug response in cancer cell lines using high-throughput gene expression data from RNA-seq. Because cancer cell lines serve as models for patient tumors, accurate predictions of drug response can be used to improve understanding of cancer systems biology and potentially inform patient treatment recommendations. Gene expression and drug response data was obtained from DepMap^26^. There are two key challenges for this dataset: (1) there are ~50,000 gene expression features but only ~1,000 cancer cell lines and ~700 drugs, so preventing overfitting is essential, and (2) the dataset is highly imbalanced because there is a small number of cell lines that respond to each drug.

Using Galaxy-ML, we built a meta-ensemble as well as other learners for each drug. The meta-ensemble included a linear boosted model, tree boosted model, and k-nearest neighbor regression, and we use principal component analysis (PCA) for dimensionality reduction in several learners. Dimensionality reduction was used in an effort to address the challenge of using a dataset with a very large number of features. We developed predictors for both regression and classification; labels for classification were generated by thresholding drug response values and labeling cell lines as responders or non-responders to each drug using a cutoff of z-score < −1 for responders. Predictors were scored using average precision to address the challenge of assessing model performance on a highly imbalanced dataset, where the goal is to identify responders (true positives) amongst a very large number of non-responders. To compare regressors and classifiers, average precision for regressors was calculated using rank-ordered predictions, which has been done in past machine learning work in this space^6^. We evaluated each learner using nested CV, with 5-fold CV for 4 repetitions for the outer splits and 5-fold CV with two repetitions for the inner splits. Our results show that stacking regressors performed best for both regression (Figure 2c) and classification (Figure 2d). Linear boosting approaches also performed very well, with results that were on par with the meta-ensembles. Successful completion of these two use cases shows that Galaxy-ML can support large and diverse machine learning experiments.

In the third use case, Galaxy-ML was used to reproduce key results from Selene^12^, a deep learning toolkit for biological sequence data built on the PyTorch library. Using Galaxy-ML, we reimplemented two deep learning architectures originally implemented in Selene that model and predict regulatory elements, including transcription factor binding sites, DNase I hypersensitive sites, and histone marks. Results from these models are within 1% of those reported for Selene (Figure 3, and Methods Section M.3). Critical to this work was the implementation of data generators in Galaxy. Data generators meet two important needs: (1) producing new examples from existing data to increase the number of instances available for training, and (2) feeding small sets of examples to the deep learning model so that the entire training dataset does not need to be loaded into memory. This use case demonstrates that Galaxy-ML deep learning tools are general and powerful enough to support realistic use cases and that Selene results are reproducible across different deep learning implementations.

**Figure 3.**
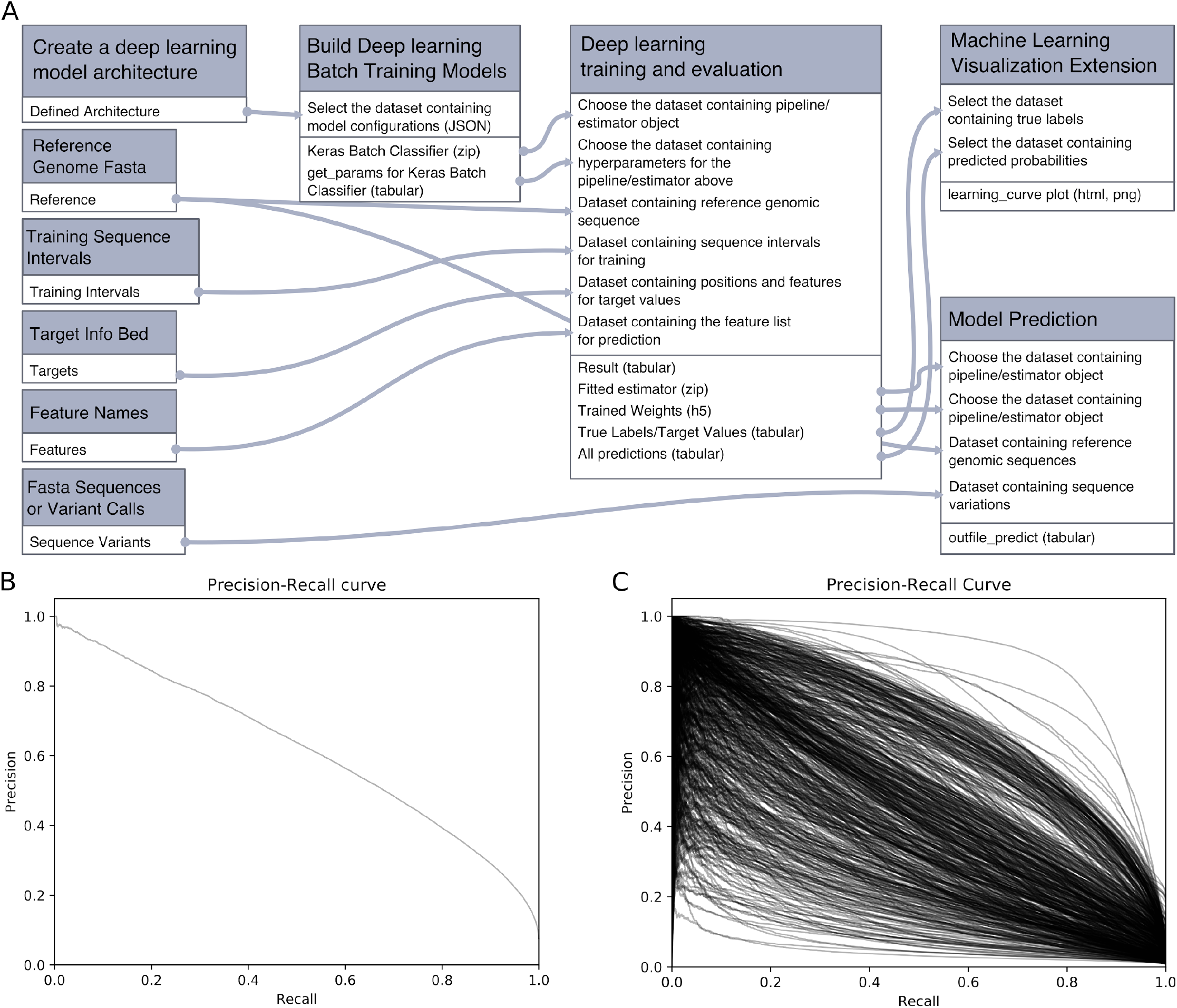
(A) Galaxy workflow to create and train a deep learning model, then use the model for visualization and prediction. (B) Precision-recall curve for a deep neural network trained to predict binding sites for a single transcription factor. (C) Precision-recall curves for a deep neural network that predicts 919 regulatory element binding profiles.

In summary, Galaxy-ML helps accelerate biomedical research by making machine learning more accessible, scalable, and reproducible for all biomedical scientists. Galaxy-ML’s tools are completely generalizable and have applications well beyond these use cases. With Galaxy’s web-based user interface, an entire machine learning pipeline from normalization, feature selection, model definition, hyperparameter optimization, and cross-fold evaluation can be created and run on a compute cluster using a web browser. This makes scalable and reproducible machine learning accessible to biomedical scientists regardless of their informatics skills. By leveraging the more than 7800 analysis tools available in Galaxy, comprehensive end-to-end analyses can be performed, which begins with primary analysis of -omics, imaging, or other large biomedical dataset and continues to downstream machine learning tools that build and evaluate predictive machine learning models from features extracted from the primary data. The website https://galaxyproject.org/community/machine-learning/ provides a hub for machine learning in Galaxy and access to all Galaxy-ML tools, workflows and tutorials. We anticipate that this hub will serve as a community starting point to foster accessible machine learning in biomedicine.

## Methods

The Galaxy tool wrappers for our machine learning suite are available at the following URLs: (1) main tools: https://github.com/bgruening/galaxytools/tree/master/tools/sklearn and (2) utilities and custom classifiers: https://github.com/goeckslab/Galaxy-ML, and the entire suite can be installed onto any Galaxy server through the Galaxy ToolShed at http://bit.ly/galaxy-ml-toolshed.

Three use-cases are discussed in the parent manuscript: (1) PennML (see M.1), (2) DepMap Cancer Cell Lines (see M.2), and (3) deep learning for genomics using Selene (see M.3). For each of the use-cases we provide links to (a) publicly accessible input datasets, (b) Galaxy workflows to run the use-case, and (c) a Galaxy history that contains our results of running the workflow on the input datasets.

### M. 1 Use Case 1: PennML benchmark

For this benchmark we trained and evaluated a variety of supervised machine learning pipelines using 164 classification datasets with both binary and multi-class classifications, and 112 datasets with real-values targets from the Penn Machine Learning Benchmark repository^25^ (Link: https://github.com/EpistasisLab/penn-ml-benchmarks). Table M.1 provides links to Galaxy workflows and histories created, and all analyses can be precisely reproduced. The Penn ML benchmark analysis in Galaxy has two parts—classification and regression.

For classification, two Galaxy histories are shared: (i) classification performance of 15 classifiers with default parameter values on 164 datasets and (ii) the same classification analysis but with optimised parameters. In this analysis, XGBoost classifier outperforms other classifiers (Figure 2) by obtaining the best performance on most of the 164 datasets. Links to the workflows for XGBoost classifier with the default and optimised parameters are listed in Table M.1. For example the workflow for XGBoost classifier to achieve the best performance has two steps:

1. Preprocessing: scales all the datasets using scikit-learn’s RobustScaler.
2. SearchCV: optimises the hyperparameters such as *number of estimators, learning rate,* and *max_depth* of XGBoost classifier.

For regression analysis, we created one Galaxy history to measure the performance of 14 regressors with the default and optimised parameters on 112 datasets. Table M.1 lists this history and associated workflows. Similar to our classification results, the XGBoost regressor records the best performance (see Figure M.1). For example the XGBoost regressor workflow achieving the best performance has one step (named SearchCV) that optimises the hyperparameters such as *number of estimators, booster* and *max_depth* of XGBoost regressor. For each hyperparameter, a range of values is specified and using grid search, all parameter combinations are tried, and performance is reported with the optimal parameter settings. All the resulting datasets from running regression algorithms with and without parameter optimization is available from: https://usegalaxy.eu/u/kumara/h/pmlbregressionanalysisjune2020. Section M.1.1 has more discussion of our regression analyses.

**Table M.1.**
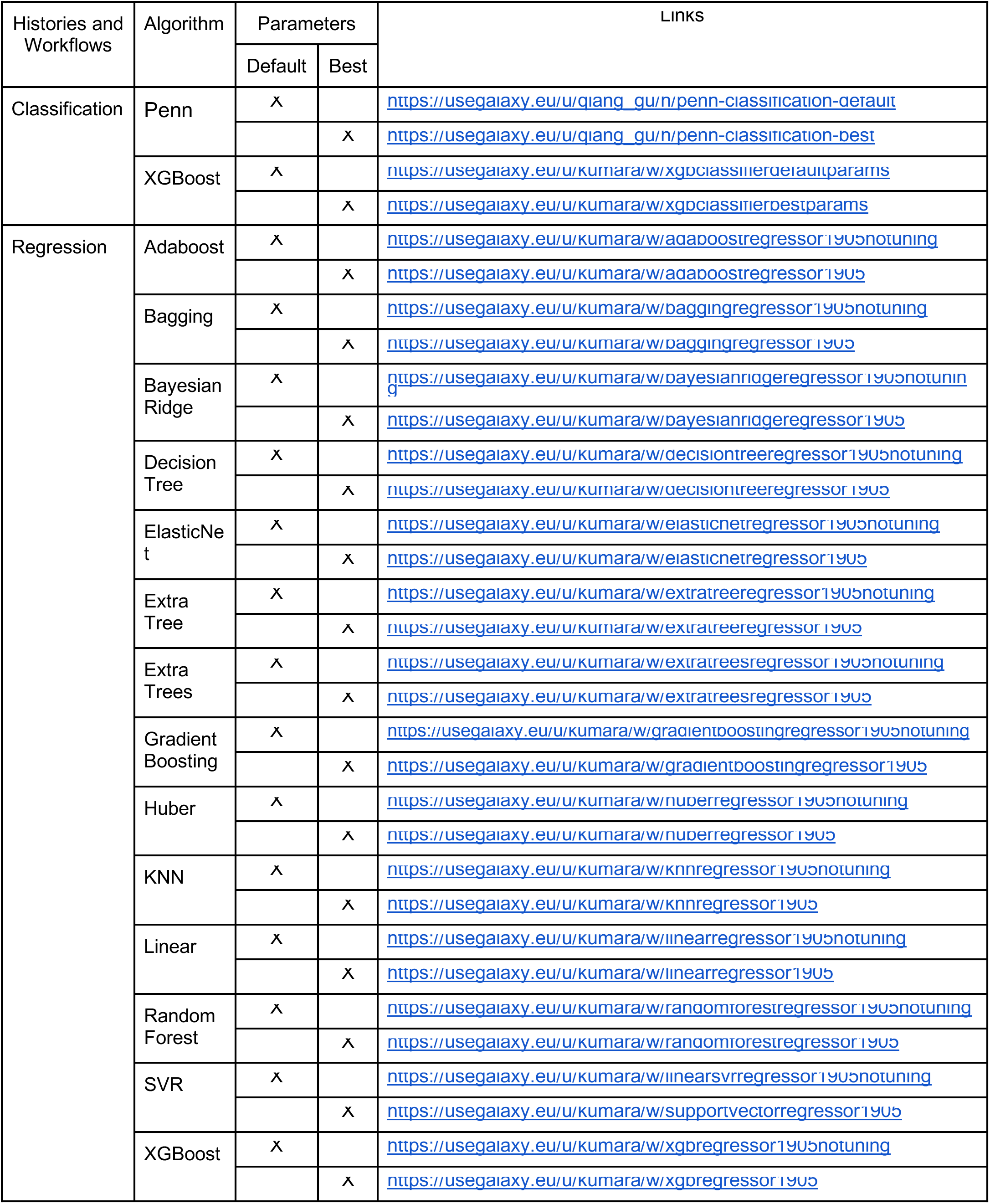
A list of Galaxy histories and workflows used for the benchmarks discussed in sections M.1 Each history/workflow ensures that an analysis can be completely reproduced because it lists all analysis steps and parameters. Each algorithm runs with two parameter configurations: default and best. Default configuration is a default value of parameters in Galaxy toolbox and best parameters are obtained by hyperparameter optimization.

#### M.1.1 Regression analysis: Comparison of 14 regressors on 112 Penn regression datasets

Using tools for data preprocessing, feature selection and regressors, we performed an aggregated analysis on 112 regression datasets from Penn Machine Learning Benchmark repository. This repository contains numerous datasets for regression many of which are of biological importance. We applied 14 different regressors on 112 datasets from the collection and performed a detailed comparison of performances of these regressors (Figure M.1). To measure the accuracy of regression models, we used the r-squared metric (R2), which is common in regression analyses. This metric can be any real number to a maximum of 1.0. If it is negative, it suggests that the regression model is not good. If it is closer to 1.0, the model’s performance is good. We used 5-fold cross-validation for training and repeated it for 10 experiment runs to compute a mean r-squared score for each dataset. We achieved a r-squared score of more than 0.80 for 3 regressors (xgboost, gradient boosting and extra trees) and close to 0.80 for 2 regressors (bagging and random forest).

**Figure M.1.**
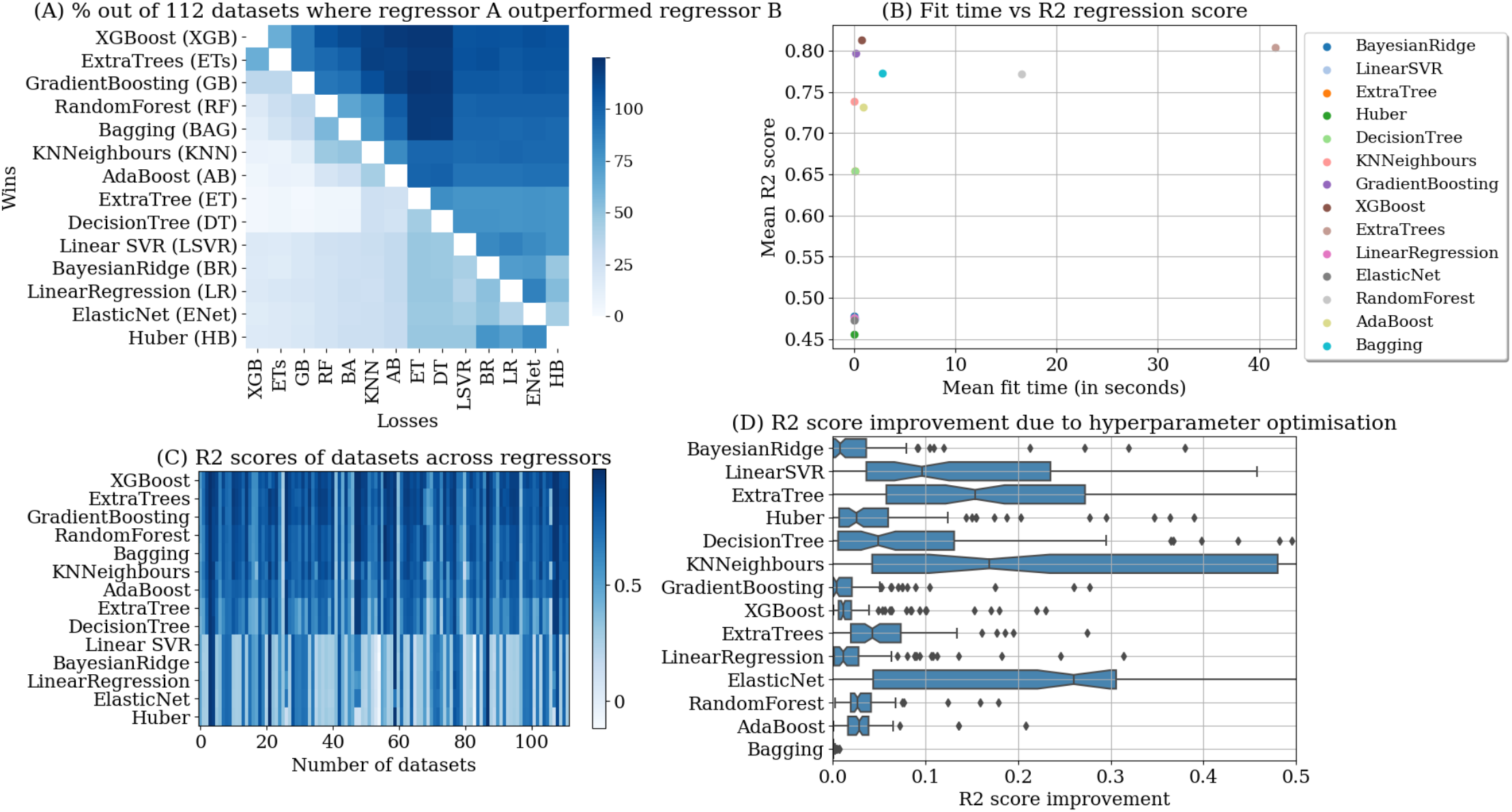
Comparison of different regression models. In panels A and C, heatmaps show the percentage of datasets for which the model listed along the row outperforms the model along the column.

Figure M.1 panel A shows a heatmap in which each square contains a number of datasets for which the regressor on the left (wins) performed better than the regressor on the bottom (losses). For example, by mapping the color of the square between adaboost (shown on y-axis) and linear regression (LR) shown on x-axis to the adjacent color-scale, we conclude that the adaboost regressor performs better on 75-80 datasets (out of 112) than the linear regressor. The subplot also shows a comparison of different regressors (on y-axis). The ensemble regressors perform better on average than the other categories which include linear, tree and nearest neighbors regressors.

Figure M.1 panel B shows a comparison between the running time and accuracy of different regressors. We compute an average running time of each regressor over all 112 datasets. The running time of a regressor on a dataset is the sum of the training and validation times for the best regression model. The regressors such as xgboost, gradient boosting and extra trees achieve > 0.80 r-squared score, but extra trees regressor requires significantly more time to finish compared to the other two regressors. Regressors such as linear regression, huber and elastic net are fast, but their accuracy is low. Decision and extra tree regressors are also fast, but their accuracy is better (> 0.7 r-squared score) than the linear regressors.

Figure M.1 (C) shows the r-squared scores of each regressor for all datasets. The linear regressors at the bottom-left of the subplot achieve lower scores than the ensemble regressors such as xgboost, gradient boosting at top-left of the subplot. We can also see that for a few datasets, none of the regressors perform well.

Figure M.1 (D) shows the importance of tuning the hyperparameters of the regressors for each dataset. It is not recommended to compute the performance of predictive algorithms over multiple datasets using the same or default values of their hyperparameters. The performance of a regressor varies for different values of hyperparameter for a dataset. Therefore, we computed the best set of values of hyperparameters for each dataset using an exhaustive search strategy (grid-search). The figure shows an improvement in r-squared scores for each regressor due to hyperparameter optimisation. Regressors such as elastic net, k nearest neighbours, decision tree and linear svr show higher improvements than bagging, random forest, adaboost, gradient boosting, xgboost, extra trees, linear regression, huber and gradient boosting in their respective r-squared scores averaged over all 112 datasets.

### M.2 Use Case 2: DepMap Cancer Cell Lines

In our second analysis, we analyzed cancer cell lines gene expression and drug response datasets from the Cancer Dependency Map Project^26^ (https://depmap.org/). This dataset includes more than 50,000 gene expression values for over 1000 cancer cell lines obtained from bulk RNA-seq as well as drug response data for 265 drugs. Both gene expression data and drug response targets are continuous data. We had several goals in mind when performing this analysis. We wanted to assess how well supervised learning performed on a dataset with a very large number of features and a relatively small and imbalanced number of examples. These challenges are common when machine learning is applied to molecular datasets. We also wanted to compare performance of meta-ensemble (stacking) approaches with traditional single-model methods.

- Galaxy History URLs:
  ○ Regression: https://usegalaxy.eu/u/qiang_gu/h/depmap-regression
    ■ Example workflow: https://usegalaxy.eu/u/kumara/w/stackingensembleregressorpcaknr
  ○ Classification: https://usegalaxy.eu/u/qiang_gu/h/depmap-classification
    ■ Example workflow: https://usegalaxy.eu/u/kumara/w/stackingclassifierdrugprna2

Because target values for this analysis—cell line drug response data—was continuous, we developed a strategy to binarize the data so that classification approaches could be used. Drug response values were standardized/z-scored, and:

- cell lines with a standardized value of less than −1 were labeled responders;
- cell lines with a standardized value between −1 and 0 wer e labeled indeterminate;
- cell lines with a standardized value greater than 0 were labeled nonresponders.

We implemented this strategy in two tools in the Galaxy-ML utils library, `binarize_average_precision_scorer` and `binarize_auc_scorer`, which report average precision and ROC AUC scores, respectively.

We compared eight regression approaches and 11 classification approaches for this dataset. Some approaches used principal component analysis to reduce the number of features before training a model. Several approaches used meta-ensembles or stacking, and grid search was used to optimize hyperparameters for the meta-ensembles. Figure 2C and 2D summarizes the results of this analysis, and Table M.1 includes histories for both the classification and regression analyses.

### M.3 Use Case 3: Deep Learning for Genomics using Selene

Using Galaxy-ML, we reproduced two deep learning models originally implemented in Selene, a deep learning library for biological sequence data^12^. The objective of the first analysis was to train an existing deep learning architecture using a novel dataset. Specifically, train the DeepSEA^27^ architecture to model a tissue-specific regulatory element for a single transcription factor not supported by DeepSEA (see https://github.com/FunctionLab/selene/tree/master/manuscript/case1). To reproduce this analysis, we used the DeepSEA architecture (https://github.com/FunctionLab/selene/blob/master/models/deepsea.py), which contains 3 convolutional layers, 2 pooling layers, one fully connected layer and a Sigmoid output layer, plus the dataset used by Selene. Both Galaxy-ML and Selene obtained the same results: a ROC AUC of 0.94 (Figure M.2 and Table M.2). The Galaxy history for this analysis used GPUs for training and evaluation, and it is available at https://usegalaxy.eu/u/khanteymoori/h/sequencedlselenecase1v080522

Another experiment run using Selene compared the performance of the DeepSEA architecture with an extended architecture that includes three additional convolutional layers (see https://github.com/FunctionLab/selene/blob/master/manuscript/case2/README.md). We reimplemented the extended architecture and then trained and evaluated this model using the Selene dataset (Figure M.3), which included data for 919 regulatory elements. Our results are nearly identical to those obtained by Selene (Table M.2). The Galaxy histories for this use-case are available from the following links:

- https://figshare.com/articles/Galaxy-History-selene-case2_tar_gz/12398699 — fully trained over 100 epochs using a private Galaxy server with acces to a GPU cluster;
- https://usegalaxy.eu/u/qiang_gu/h/selene2 — online Galaxy history trained for only two epochs using CPU cluster.

**Figure M.2.**
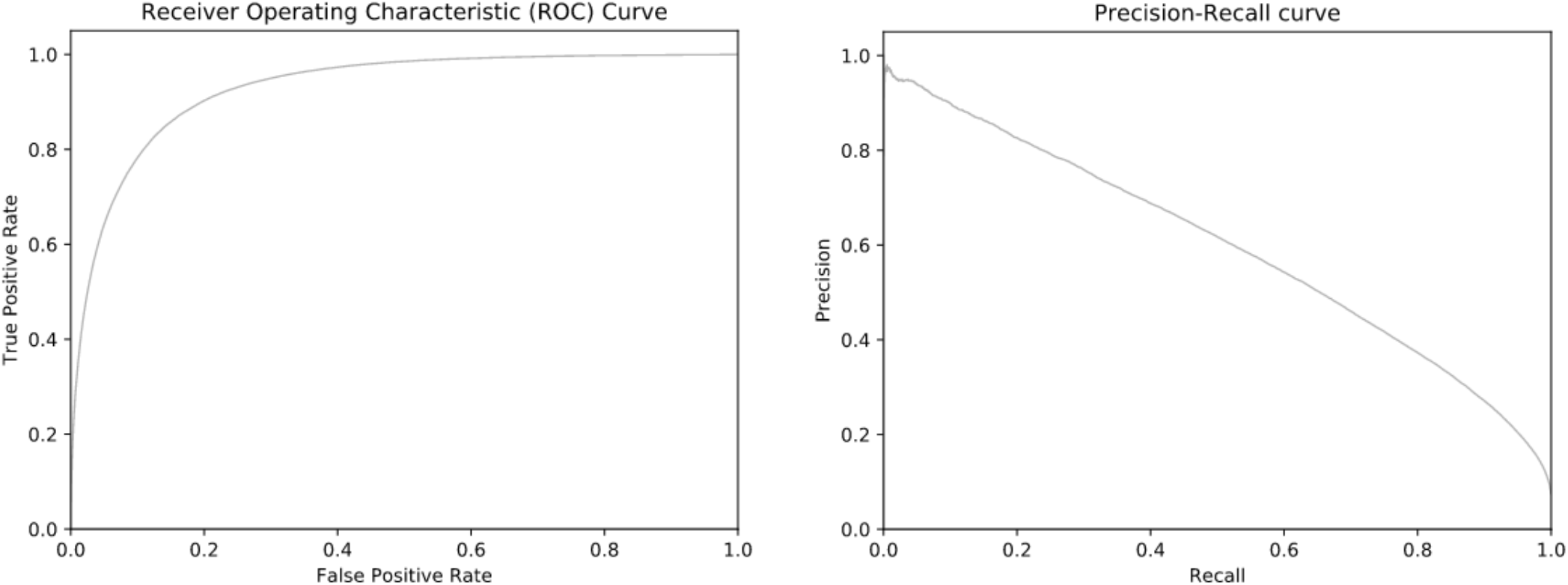
Visualized results obtained using the DeepSEA architecture to model regulatory elements for a single tissue-specific transcription factor.

**Figure M.3.**
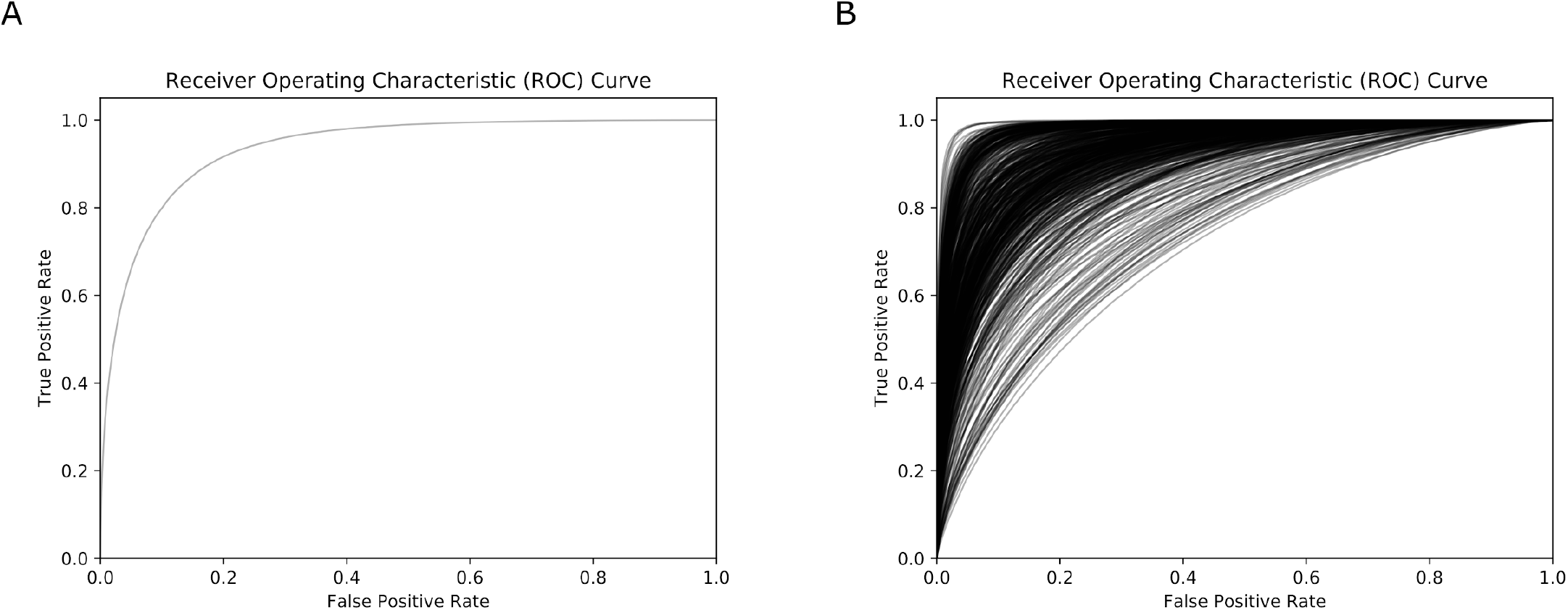
Visualized ROC curve results obtained from using the extended DeepSEA architecture to model 919 regulatory elements: (A) average ROC for all elements and (B) individual ROC curves for each element.

**Table M.2.** Performance results obtained using Galaxy-ML models fully trained using GPU and Selene models. All datasets used were obtained from Selene. AUPRC is the area under the precision-recall curve, and is also known as the average precision. “N.R.” means that the models did not report this information.

| Analysis/<br>Case Study | Galaxy-ML |  |  | Selene |  |
| --- | --- | --- | --- | --- | --- |
|  | Balanced Accuracy | ROC AUC | AUPRC | ROC AUC | AUPRC |
| 1 | 0.68 | 0.94 | 0.62 | 0.94 | N.R. |
| 2 | 0.62 | 0.928 | 0.365 | 0.938 | 0.362 |

